# Role of vegetation characteristics on the distribution of three hornbill species in and around Pakke Tiger Reserve, Arunachal Pradesh, India

**DOI:** 10.1101/2022.01.01.474716

**Authors:** Soumya Dasgupta, Tapajit Bhattacharya, Rahul Kaul

**Affiliations:** Wildlife Trust of India, F 13, Sector 8, Noida, Uttar Pradesh - 201301; Wildlife Institute of India, Chadrabani, Dehradun - 248001; Post graduate department of Conservation Biology, Durgapur Government College, Jawahar Lal Nehru Road, Amarabati Colony, Durgapur, West Bengal 713214

**Keywords:** Hornbill abundance, North East India, forest structure, mutualism

## Abstract

The relationship between various vegetation characteristics and the relative abundance of three hornbill species [Great Pied Hornbill (*Buceros bicornis*), Wreathed Hornbill (*Rhyticeros undulatus*) and Oriental Pied Hornbill (*Anthracoceros albirostris*)] was studied in and around Pakke Tiger Reserve, Arunachal Pradesh. We walked transects (n=11; 22 walks) in three study sites to detect hornbills. Vegetation sampling was done using circular plots (n=33; 10 m radius) at every 400m interval along each transect. Encounter rate (1.5± 0.188/km) of Great Pied Hornbill (*Buceros bicornis*) was highest in the protected and undisturbed forest area where food and roosting tree density were also high (114/ha). Oriental Pied Hornbill was common in both the sites within Pakke Tiger reserve near riverine forests (0.75±0.25/km) and also in the dense undisturbed forest (0.875±0.226/km). Multivariate analysis revealed that tree density, presence of fruiting trees (utilized by hornbills), canopy cover, and tree diversity in a particular area are the major factors responsible for the assemblage of more than one species of hornbills. The study shows that protection of the forest patches to keep the diversity and density of the tree species intact is crucial for the survival and distribution of the hornbills in the landscape.

## Introduction

In the tropical rainforest, plant-frugivore interaction is important for the sustenance of the diversity of the plant species. Most tropical species depend on vertebrate frugivores for the dispersal of seeds^1, 2, 3^. Seed dispersal increases the chance of seed survival by decreasing the density-dependent mortality of the seeds under the own tree canopy and increasing the fitness of individual plants by influencing the distribution of plant species spatially in a large landscape^4, 5^. Studies in South-East Asia shows that small assemblages of potential dispersers disperse large seeds ^6,7^. Large bodied frugivores with larger mouth capacity and gape widths can feed on a wide range of fruits with different sizes^8, 3^ and thus can potentially disperse seeds of larger sizes, which can’t be dispersed by animals with smaller gape size or mouth cavity ^2, 9^. Since dispersal by vertebrates is the most common dispersal mode of the 50-90% of the tropical trees^10,11^, a reduced abundance of frugivores could affect the dispersal, regeneration, and eventually the demographics of the dependent tree species ^2, 6^.

In a mutualistic manner, vegetation is a major determining factor of species assemblages in a bird community and the abundance of certain bird species depends on the vegetation composition of the habitat ^12, 13, 14,15, 16^. Both physiognomic and floristic characteristics of vegetation affect the bird species abundance^16, 17^. Change in vegetation along the complex environmental and disturbance gradient can change the abundance and distribution of a particular species or species guild ^18^ and the interplay between vegetation and bird species assemblage was well documented for decades ^18,19,20, 21,22^. Bird species richness and diversity are directly correlated with the physiognomic characters such as foliage height, foliage volume, percentage vegetation cover, etc. ^23, 24, 25, 20^. The abundance of some tree species or a group of species can influence the distribution of certain bird species or bird guilds ^26, 27^. Large bodied bird species such as hornbills are mutualistic in the forest. Regeneration of forest vegetation depends a lot on the seed dispersers (such as hornbill) presence in the forest ^28^. Seed dispersal in the tropical forest is one of the vital ecosystem services ^29, 30, 31^, and in the absence of dispersers, loss of diversity is well observed throughout the world^6^.

Seed dispersers are particularly sensitive to both habitat destruction and hunting ^32, 33, 34, 35, 36^. A considerable amount of habitat alteration and loss has intensified in the tropical forest of North East India in the Eastern Himalayan region ^37, 38^. Hunting, loss of habitat due to jhum (shifting cultivation), and developmental activities are severe in the easily accessible foothill forests near the dense human habitation^36^. Remaining forest patches are also under threat due to rampant hunting and extraction of forest products ^39, 40, 41^. Pakke Tiger Reserve in Arunachal Pradesh is an apt example of such habitat alterations. Baseline information on the hornbill abundance and its role in seed dispersal is available from different forests of Arunachal Pradesh, including Pakke Tiger Reserve ^6,34,36,22^. From Pakke Tiger Reserve, Datta and Rawat^42^ (2008) indicated that tree species of Lauraceae, Meliaceae, Myristicaceae, Anacardiaceae, Clusiaceae, Burseraceae, Moraceae, and Euphorbiaceae are used for fruit. Seed dispersal by animals and ornithochory were the most abundant dispersal modes of tree species in Pakke Tiger Reserve ^42, 43^.

However, information on the relationship between hornbill species occurrence and vegetation/stand characteristics are still lacking in this landscape. The present study was carried out with two primary objectives, first to assess the occurrences and relative abundances of different hornbill species in and around the Eastern boundary of Pakke Tiger Reserve. Second, how both physiognomic and floristic characteristics of vegetation determine the relative abundances of different hornbill species.

## Methods

### Study area

The study was done in the lowland tropical forest of Pakke Tiger reserve in Arunachal Pradesh. We selected three study sites, two within Pakke Tiger Reserve (*West Bank* 26° 56′ N, 92° 58′E and *Khari* 26°59′ N, 92° 54′E) and one outside the Park boundary (Lanka 27°01′ N, 93°02′E) (Figure 1). The major vegetation type of the study area is classified as Assam Valley tropical semi-evergreen forest (2B/C1) by Champion and Seth (1969)^44^. Details of different habitat types and levels of disturbance can be found in Datta (1998) ^34^. All three sites were selected based on the forest structure and degree of disturbances. *West Bank* forest (12 Km^2^) is situated near *Seijusa* on the southern boundary of the Park adjacent to the Pakke River and is free of large-scale anthropic interventions or habitat alteration. The area is dominated by lowland forests with an altitudinal range of 150m to 600m. *Khari* is located almost 12 km west of West Bank and is characterized by steep terrain dominated by palm (*Livistona*), cane (*Calamus*), and bamboo (*Bambusa* and *Dendrocalamus*) species, intersected by several perennial streams. Human settlements practicing log and cane extraction existed here in the past (2-3decades ago)^34^. *Lanka*, on the other hand, is outside the boundary of Pakke Tiger Reserve and therefore is heavily disturbed due to the sustained prevalence of selective logging and hunting. Local people also heavily depend on the forests of *Lanka* for firewood and other non-timber forest products.

**Figure 1:**
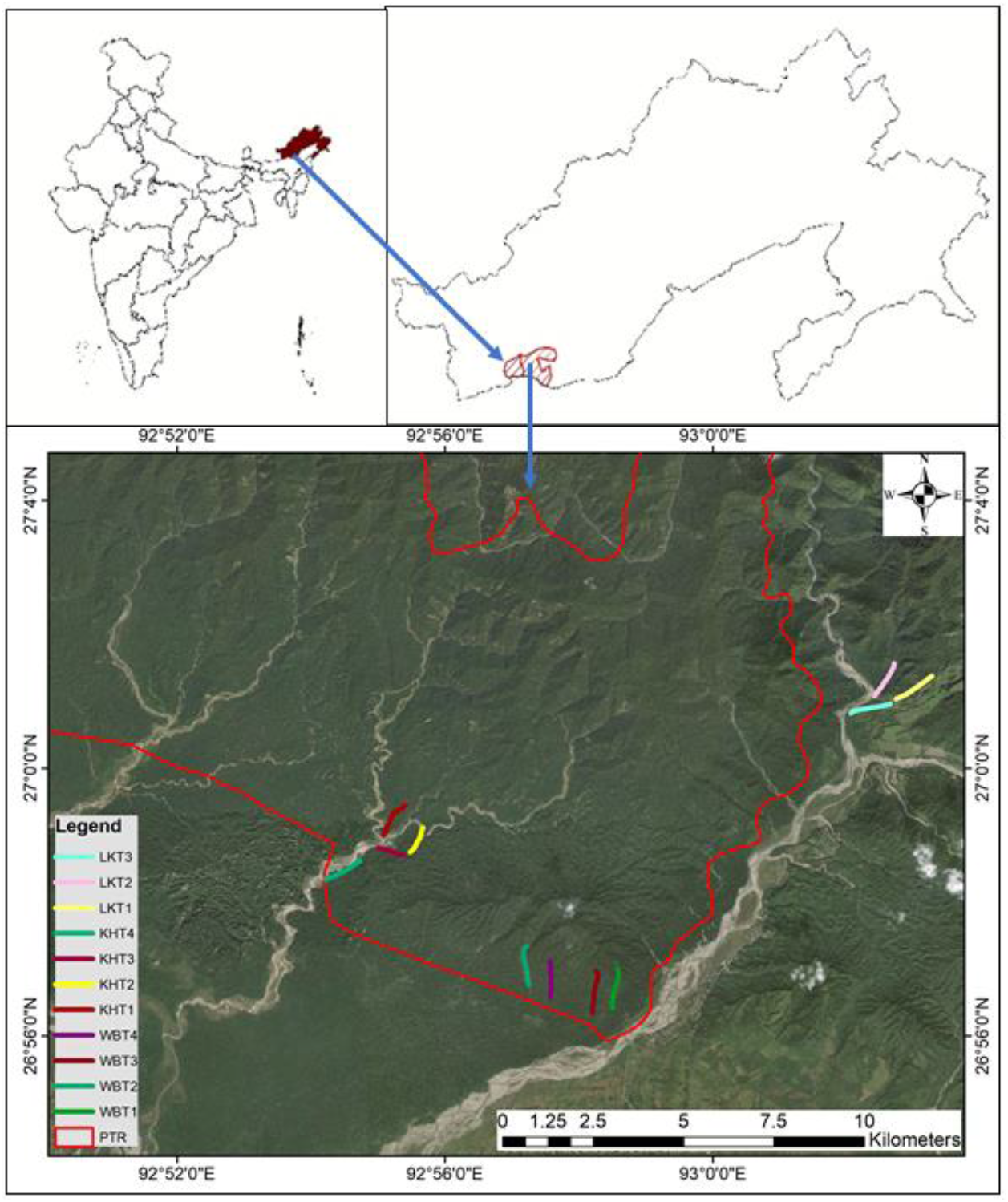
Map of the Study area in Pakke Tiger Reserve, Legends showing the Transects in Lanka (LK) khari (KH) and West Bank (WB) and Boundary of the Pakke Tiger Reserve

### Field methods

We walked transect (n=11; 22 walks) of one km each in three study sites to detect hornbill species from 5.00 to 7.00 hrs. As hornbill activity is high during the period and the chance of an encounter is also very high. Four transects each in *West Bank* and *Khari* and three in *Lanka* were walked twice, thus the total distance walked was 22 km, eight km each in *West Bank* and *Khari* and six km in *Lanka*. The relative abundance of hornbill species was calculated using the encounter rate per kilometer walk. Vegetation sampling was carried out using circular plots (n=33; 10 m radius) at every 400m interval along each transect (*i*.*e*. three vegetation plots in each transect). In each plot, the presence of tree (>20 cm girth), presence of climber, trail, number of fruiting, nesting, and roosting trees were recorded. Girth at Breast Height (GBH) for each tree inside the plot was measured using standard methodology^45^. The number of trees having more than 20cm GBH was identified and their height was measured. Canopy cover in each plot was measured using line intercept method^46, 47^.

### Analysis

As the number of encounters was not enough to obtain statistically reliable results using the DISTANCE software ^48^ (Thomas et al 2010), for calculation of Encounter rates, we used the following formula:

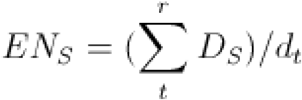

Where ‘S’ is a particular hornbill species, ‘D’ is the detection of the particular species ‘S’, and’ denotes the total distance of a particular transect ‘t’, in region ‘r’ and EN is the encounter rate. Encounter rates across individual transects were then averaged for all replications, for each study area and individual hornbill species, to arrive at relative abundance estimates per site. The standard error of means was calculated to gauge the variation across averaged encounter rates for each transect.

Identification of trees, shrubs, and herbs were carried out following Datta (2001) ^49^, and species inventory was prepared for the three study sites. Tree density, diversity, dominance, average height, and average GBH, were calculated for each vegetation plot. Relative density, relative frequency, and relative dominance of important fruiting and roosting trees for hornbills were summed to calculate the Importance Value Indices (IVI) in three different sites. Important fruiting and roosting trees of the area were already mentioned by Datta (2001)^49^.

Permanova test^50^ was done to assess the difference in plant community composition between the three study sites. Simper analysis^51^ was done to know the species influencing the difference in the community structure of the three study area with different disturbance history. The change in abundance of different hornbill species along the difference in floristic characters was assessed through non-metric multidimensional scaling (NMDS)^52^ with Bray Curtis distance measures. The abundance of different plant species with IVI values more than five was used for ordination along with the presence data of the hornbill. The importance of physiognomic variables was assessed through binomial regression. Using physiognomic characteristics as explanatory variables Generalized Linear Modeling was carried out to identify the factors governing the occurrence of different hornbill species using binomial family in R software^53^. The information generated from 33 vegetation plots were used along with the presence-absence information of hornbill species within the 300 meter stretch of the transect in both sides of the vegetation plot (Table 1). We used corrected Akaike Information Criterion^54^ (AIC_c_) values to rank the models and considered all the models whose ΔAIC < 2 as equivalent. The summed model weight of each explanatory variable in these models was used to determine the most influential variables for each species. The value and sign of the logistic coefficient of each variable (positive or negative) and the associated error was used to determine the direction of influence of the variable.

**Table 1.** List of Variables, their abbreviations and range as used to determine the influential vegetation characteristics and their role in hornbill distribution using Non Metric Multidimensional Scaling (NMDS) followed by Generalized Linear Modelling

| Abbreviation | Full form | Description |
| --- | --- | --- |
| GPH | Great Pied Hornbill | 0-1 |
| WH | Wreathed Hornbill | 0-1 |
| OPH | Oriental Pied Hornbill | 0-1 |
| diversity | Diversity of tree species | Diversity of tree species in the vegetation plot along the transect (2-19) |
| avght | Average height of the tree species | Average height of the tree species in the vegetation plot along the transect (9.58-17.67m) |
| avggbh | Average gbh of the tree species | Average gbh of the tree species in the vegetation plot along the transect (41.7-149.8cm) |
| density | Density of tree species | Density of tree species in the vegetation plot along the transect (5-27 trees/plot) |
| ftree | Presence of fruiting tree | Presence of fruiting tree of hornbill food preference in the vegetation plot along the transect (0-4) |
| rntree | Presence of roosting and nesting tree | Presence of roosting and nesting tree of hornbill in the vegetation plot along the transect (0-2) |
| canopy | Canopy cover | Percentage canopy cover in the vegetation plot along the transect (60-90%) |
| trail | Presence of trail | Presence of trail in the vegetation plot along the transect. (0-3) |
| climber | Presence of climber and epiphytes | Presence of climber and epiphytes in the vegetation plot along the transect (7-45) |

## Results

During transect walks we detected the presence of three hornbill species (Table 1) such as Great Pied Hornbill (*Buceros bicornis*), Wreathed Hornbill (*Aceros undulatus*), and Oriental Pied Hornbill (*Anthracoceros albirostris*). Encounter rate (1.5± 0.188/km) of Great Pied Hornbill was highest in the protected and undisturbed forest area of *West Bank* (Table 2). The encounter rates of the other two species were also higher in undisturbed protected forests than the other two sites.

**Table 2.** Average values of different physiognomic variables and presence of different hornbill species in different sites as well as transects. Hornbill presence was shown as three if it was encountered in all the three segment of the transect and decreased as the encounter decreased along the transect.

| Site/Transect | Average GBH | Average Tree Height | Average Tree Density | Canopy cover | Great Pied Hornbill | Wreathed Hornbill | Oriental Pied Hornbill |
| --- | --- | --- | --- | --- | --- | --- | --- |
| WestBank T1 | 57.17±49.65 | 12.47±6.66 | 21.6 | 81.66 | 3 | 1 | 2 |
| WestBank T2 | 52.15±37.63 | 14.46±5.82 | 22 | 85 | 2 | 2 | 1 |
| WesBankT3 | 66.6±38.78 | 15.15±4.54 | 21.6 | 81.66 | 1 | 1 | 2 |
| WestBankT4 | 58.73±36.32 | 14.66±6.03 | 18.6 | 85 | 3 | 2 | 1 |
| Khari T1 | 67.84±44.20 | 12.23±4.81 | 21.3 | 83.33 | 0 | 1 | 0 |
| Khari T2 | 67.43±46.73 | 11±4.67 | 18.3 | 81.66 | 1 | 0 | 2 |
| Khari T3 | 57.2±31.96 | 11.95±4.25 | 16 | 81.66 | 1 | 1 | 2 |
| Khari T4 | 51.03±25.98 | 11.61±3.31 | 18 | 80 | 2 | 0 | 1 |
| Lanka T1 | 69.67±38.68 | 11.05±3.99 | 13 | 80 | 1 | 1 | 1 |
| Lanka T2 | 83.35±40.23 | 14.36±5.40 | 15 | 78.33 | 1 | 0 | 2 |
| Lanka T3 | 108.2±81.03 | 12.56±4.96 | 8 | 63.33 | 1 | 2 | 0 |

From the vegetation plots of three study sites, a total of 62 tree species were recorded among which 56 were identified up to species level. When tree diversity and richness were measured, the Shannon index gave high measures in Lanka (3.32) among the three sites, followed by West Bank (3.08) and Khari (2.69). The dominance index on the other hand was much higher in Khari (0.15) than that of West Bank (0.06) and Lanka (0.04). The average GBH (girth at breast height) was higher in Lanka compared to the trees measured in West Bank and Khari (Table 2), whereas the opposite trend was found in the case of tree density in the vegetation plots. Average tree height and canopy cover range from 11 to 15.153 meter and 63.33 percent in one transect of Lanka to 85 percent in two transects of West Bank. The details of the measurements were given in Table 2. The presence-absence of three hornbill species near the vegetation plots were assessed during the data analysis. Great Pied Hornbill was found near all the three vegetation plots of two transects of West Bank, whereas it was not encountered during the transect walk in one of the transects in Khari. Wreathed Hornbill and Oriental Pied Hornbill were not encountered in one transect of Khari and Lanka each and none of the two hornbill species were encountered throughout the transects in all the three sites (details given in table 2). The result of Non-metric multidimensional scaling (NMDS) depicts that the presence of Wreathed Hornbill and Great Pied Hornbill determined by similar factors like high diversity of trees, trees with larger GBH and height and association with tree species like *Polyalthia simiarum, Chisocheton sp, Pterospermum sp, Dysoxylum sp*., etc., whereas the presence of Oriental Pied Hornbill is associated with canopy cover and tree density (Figure 2). The Importance Value Indices of *Polyalthia simiarum* and *Chisocheton sp* were higher among all the 13 important fruiting and roosting trees in both the undisturbed sites (Figure 3) of *West bank* and forests of *Khari* where human intervention occurred in the past. However, their IVIs were zero or minimal in the disturbed sites of *Lanka*.

**Figure 2.**
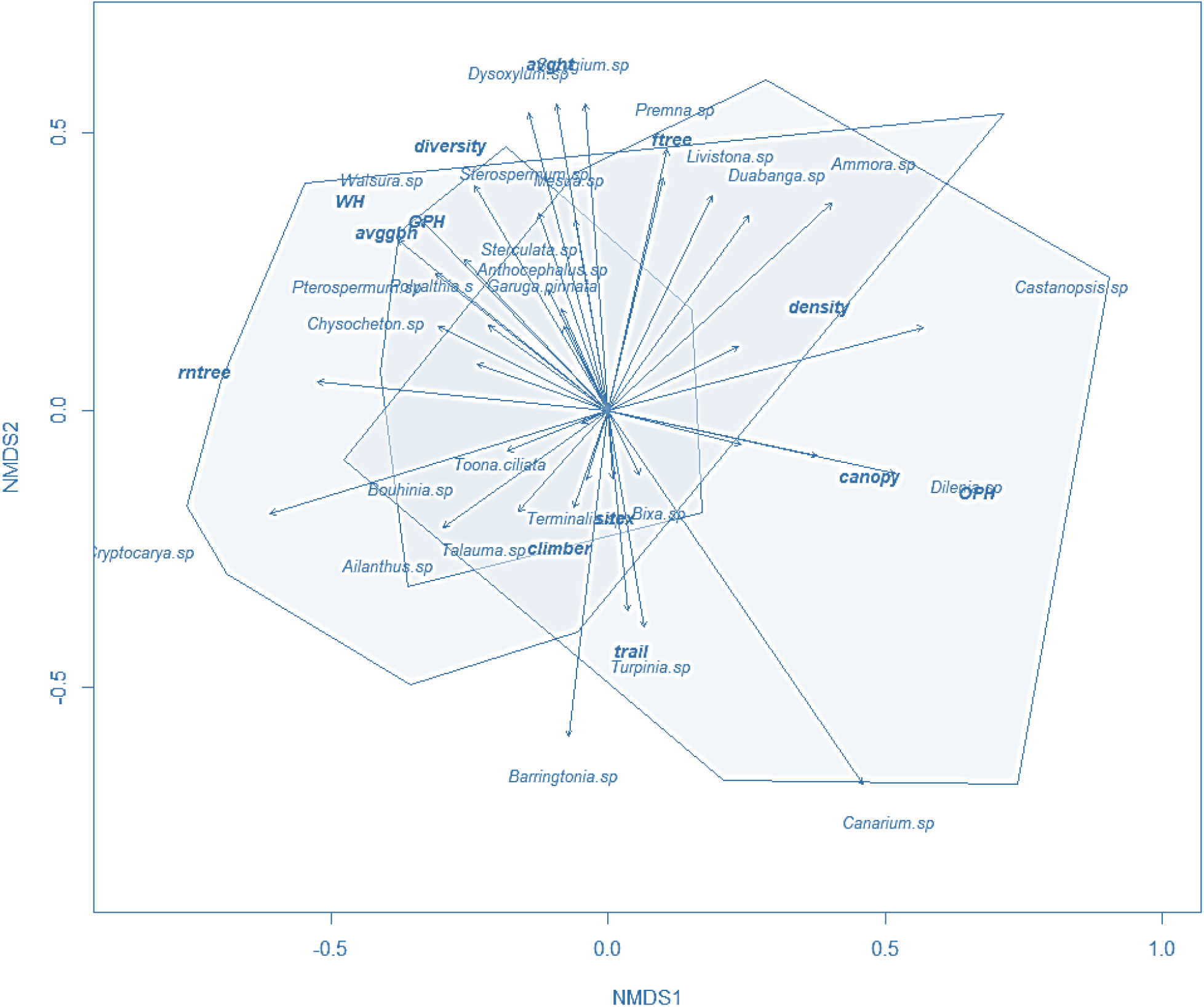
Result of Non Metric Multidimensional Scaling showing the interaction of physiognomic and floristic characteristics with the hornbill presence.

**Figure 3:**
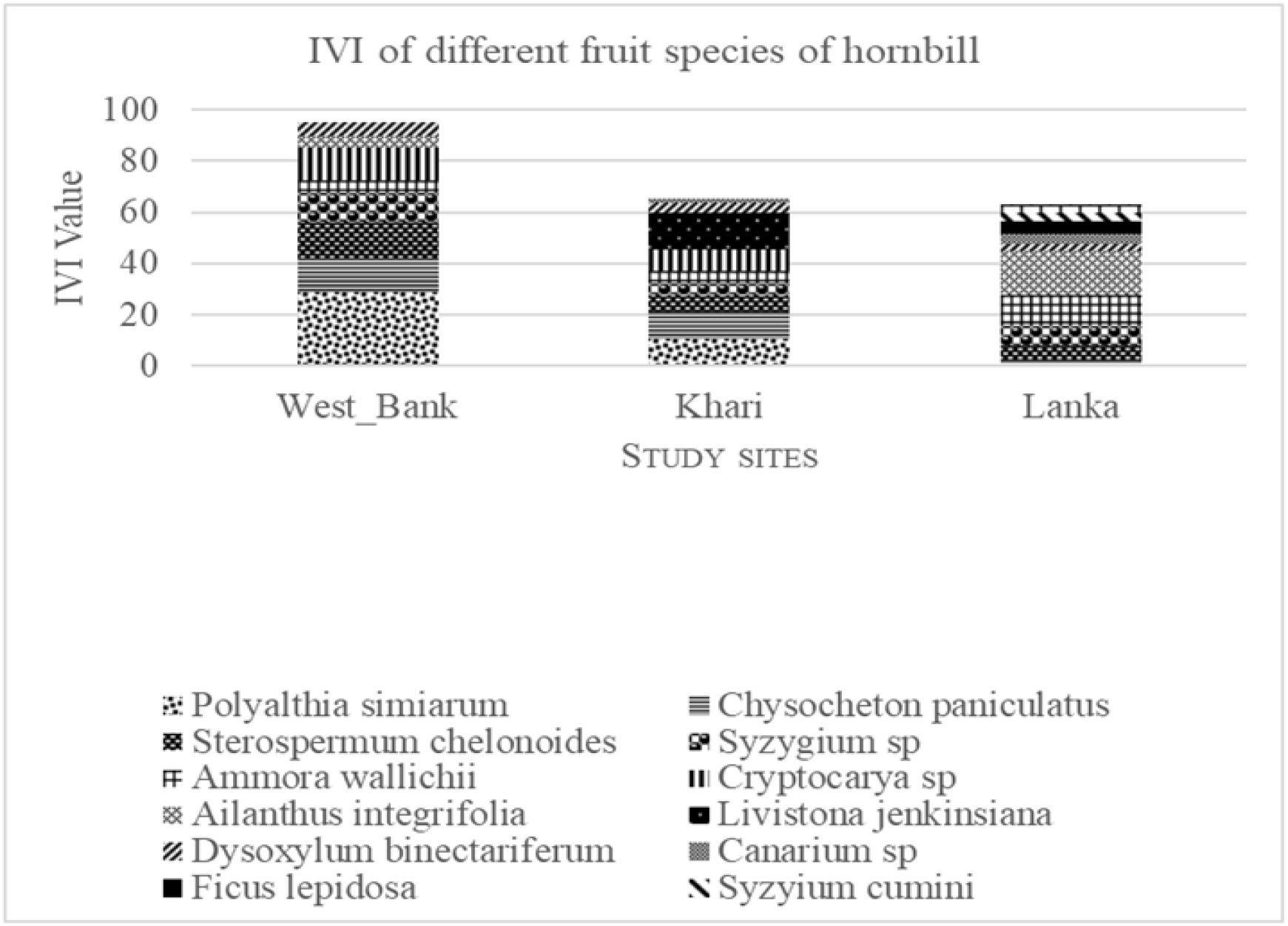
Importance value indices of different fruiting and nesting trees depicted in stacked column chart

The NMDS results also shows some differential association of species in three different study sites. Species like *Terminalia, Garuga pinnata, Anthocephalus sp, Sterculata sp* were only found in West Bank and forms a distinct association. *Pterospermum sp, Polyalthia sp* and *Chisocheton sp* forms a distinct distribution although the abundance of these species also contributing the dissimilarity between the tree community in the study area. The result of Permanova shows a significant difference (p=0.001, F= 4.32) in the species assemblage in the tree community in the three studied sites. The SIMPER analysis shows that the species assemblage of vegetation plots in Khari and Lanka differ much more (Percentage dissimilarity 93.49) than that of between West Bank and Khari (Percentage dissimilarity 89.97) and West Bank and Lanka (percentage dissimilarity 85.81). The overall dissimilarity of the tree composition is 89.78 percent in between the three community and abundance of trees of *Canarium sp, Polyalthia sp, Cryptocarya sp, Pterospermum sp* and *Bauhinia sp* were contributing more than fifty percent of the dissimilarities (Table 3).

**Table 3.** Species responsible for the average dissimilarity in between the sites and their contribution as per SIMPER analysis

| Taxon | Average.<br>dissimilarity | Contrib.<br>% | Cumulative<br>% | Mean abund.<br>In West<br>Bank | Mean abund.<br>In Khari | Mean abund.<br>In Lanka |
| --- | --- | --- | --- | --- | --- | --- |
| Canarium sp | 18.6 | 20.72 | 20.72 | 0 | 6.42 | 0.111 |
| Polyalthia s | 8.624 | 9.605 | 30.32 | 2.25 | 0.75 | 0 |
| Cryptocarya sp | 7.175 | 7.991 | 38.31 | 1.33 | 0.583 | 1 |
| Pterospermum<br>sp | 6.06 | 6.749 | 45.06 | 0.833 | 1.25 | 0.333 |
| Bouhinia sp | 5.472 | 6.094 | 51.16 | 1.25 | 0 | 0.667 |
| Castanopsis sp | 4.712 | 5.248 | 56.4 | 0.167 | 1.08 | 0.222 |
| Chisocheton sp | 4.711 | 5.247 | 61.65 | 1.08 | 0.583 | 0.111 |
| Syzygium sp | 3.544 | 3.948 | 65.6 | 0.833 | 0.25 | 0.333 |
| Sterculata sp | 3.203 | 3.567 | 69.17 | 1 | 0 | 0 |
| Sterospermum<br>sp | 2.952 | 3.289 | 72.46 | 0.667 | 0.25 | 0.222 |
| Livistona sp | 2.839 | 3.162 | 75.62 | 0 | 0.75 | 0 |
| Premna sp | 2.686 | 2.992 | 78.61 | 0.417 | 0.25 | 0.333 |
| Ailanthus sp | 2.463 | 2.744 | 81.35 | 0.167 | 0 | 0.667 |
| Talauma sp | 1.968 | 2.192 | 83.54 | 0.167 | 0 | 0.556 |
| Ammora sp | 1.727 | 1.924 | 85.47 | 0.0833 | 0.167 | 0.333 |
| Dysoxylum sp | 1.674 | 1.864 | 87.33 | 0.333 | 0.167 | 0.111 |
| Barringtonia sp | 1.595 | 1.777 | 89.11 | 0.25 | 0.0833 | 0.222 |
| Toona ciliata | 1.4 | 1.56 | 90.67 | 0.417 | 0 | 0 |
| Dilenia sp | 1.384 | 1.541 | 92.21 | 0 | 0.417 | 0 |
| Walsura sp | 1.377 | 1.534 | 93.74 | 0.0833 | 0 | 0.333 |
| Bixa sp | 1.344 | 1.497 | 95.24 | 0.417 | 0 | 0 |
| Mesua sp | 1.33 | 1.482 | 96.72 | 0.417 | 0.0833 | 0 |
| Turpinia sp | 1.033 | 1.151 | 97.87 | 0 | 0.333 | 0 |
| Duabanga sp | 0.9187 | 1.023 | 98.9 | 0 | 0 | 0.222 |
| Garuga pinnata | 0.4896 | 0.5454 | 99.44 | 0.167 | 0 | 0 |
| Terminalia sp | 0.2562 | 0.2853 | 99.73 | 0.0833 | 0 | 0 |
| Anthocephalus<br>sp | 0.2448 | 0.2727 | 100 | 0.0833 | 0 | 0 |

The selected models for presence of three hornbill species as per the lowest corrected Akaike Information Criteria (AICc) value, were given in Table 4. The result of generalized linear modeling shows presence of Great Pied Hornbill is positively associated with the presence of fruiting tree, roosting and nesting tree and negatively with that of tree density and presence of trail. Wreathed Hornbill presence is associated positively with tree diversity and average GBH, whereas negatively with tree density and presence of climber. Oriental Pied Hornbill presence is associated with the presence of fruiting tree, trail, and high canopy cover (Figure 4).

**Table 4.** Top generalized linear models explaining the roles of different vegetation characteristics on distributions of great pied, wreathed and Oriental pied hornbills in and around Pakke Tiger Reserve

| Hornbill Species | Model | Log Likelihood | AICc | ΔAICc | Weight |
| --- | --- | --- | --- | --- | --- |
| Great Pied Hornbill | ~fruiting tree | -21.053 | 46.5 | 0.00 | 0.362 |
|  | ~fruiting tree+Roosting and nesting tree+ tree density | -18.679 | 46.8 | 0.28 | 0.314 |
|  | ~fruiting tree+roosting and nesting tree | -20.372 | 47.6 | 1.07 | 0.212 |
|  | ~fruiting tree+ trail | -21.015 | 48.9 | 2.35 | 0.112 |
| Wreathed Hornbill | ~climber | -18.987 | 42.4 | 0.00 | 0.377 |
|  | ~climber +Average GBH | -18.285 | 43.4 | 1.01 | 0.226 |
|  | ~climber +average GBH+ tree diversity | -17.479 | 44.4 | 2.01 | 0.148 |
|  | ~average GBH+ tree density | -20.327 | 45.1 | 2.68 | 0.104 |
| Oriental Pied Hornbill | ~Canopy cover | -19.654 | 43.7 | 0.00 | 0.494 |
|  | ~Canopy cover+Fruiting Tree | -19.218 | 45.3 | 1.56 | 0.227 |
|  | ~Canopy cover+ Fruiting tree+tree diversity+presence of trail | -16.959 | 46.1 | 2.43 | 0.146 |
|  | ~fruiting tree | -20.961 | 46.3 | 2.61 | 0.134 |

**Figure 4.**
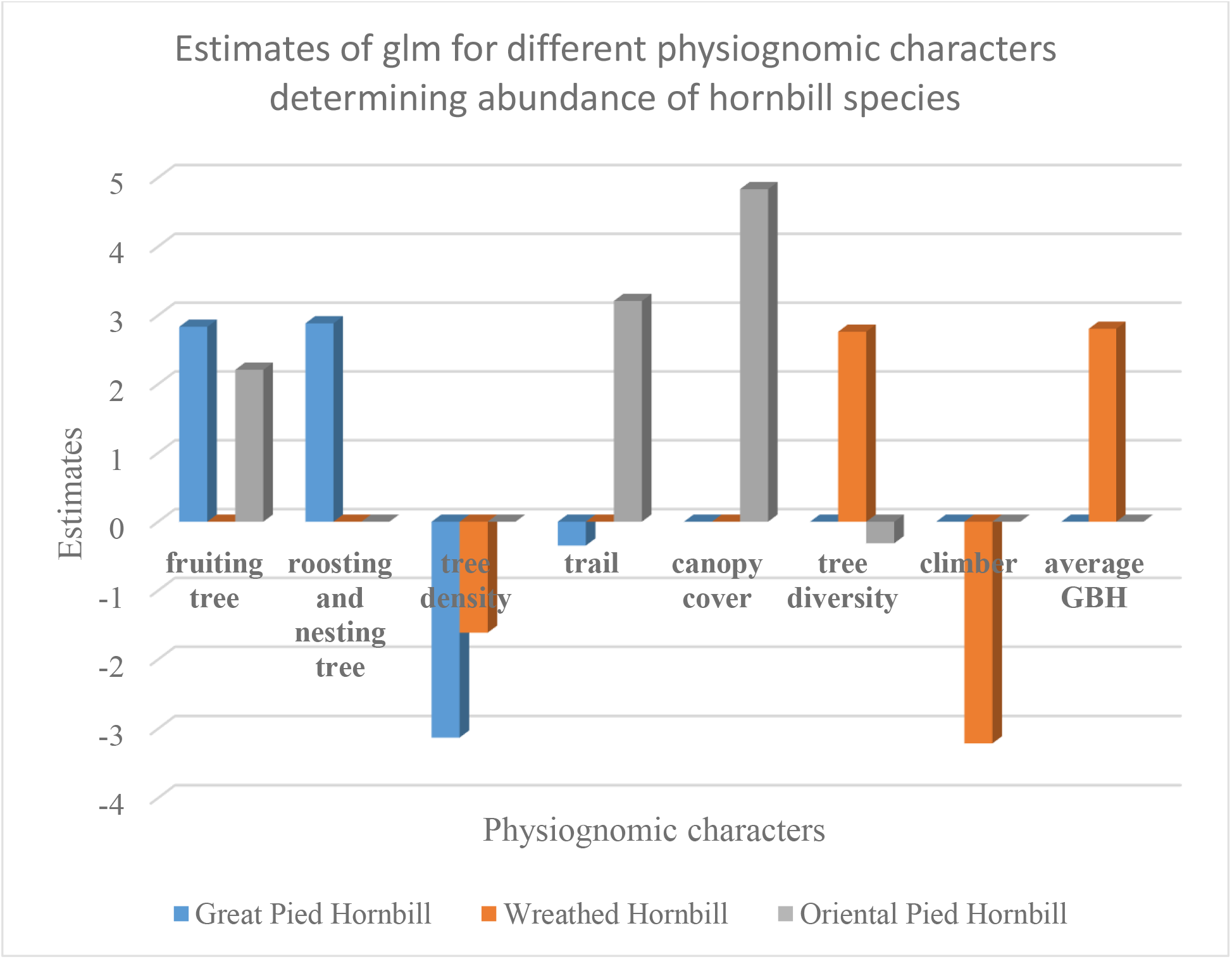
Average Estimates of the different physiognomic variables determining the hornbill presence as per the Generalized Linear Modeling (glm).

## Discussion

Previous studies in different landscapes of Arunachal Pradesh showed that the habitat preferences of these three species differ^34, 36, 49^ Great Pied Hornbills prefer undisturbed habitats, Oriental Pied Hornbills prefer riverine forest patches with secondary growth and Wreathed Hornbills show no specific habitat preferences^42^. Our current observations are comparable with these findings. Great Pied Hornbill and Wreathed Hornbill encounter rate depend on tree density, fruiting tree density, and density of roosting and nesting tree. As communal roosting is common in Great Pied and Wreathed Hornbill ^42,49^ the presence of nesting and roosting tree plays a significant role in their abundance. Whereas the abundance of Oriental Pied Hornbill also depends on high percentage canopy cover besides the presence of fruit tree and roosting and nesting tree ^34,^.

Though inside the Protected Area, low species diversity and high dominance value of the sampled plant community in *Khari* resulted from the past disturbances such as extraction of cane (*Calamus*) and bamboo (*Bambusa* and *Dendrocalamus*). *Lanka* was outside of the Protected Area and selective logging was a continuous practice, however the tree species diversity was comparable with other two sites but the tree species composition differs. In *Lanka*, the occurrence and importance value indices of trees important for hornbills differ from those of the other two sites inside the Protected Area. The change in species composition is well observed in the analysis of tree species assemblage in the three study sites. There were difference in overall species presence in *Westbank, Khari* and *Lanka*. These changes in structure and composition of tree community can explain the different species assemblage patterns of the three hornbill species as observed in this study.

Encounter Rates of all hornbill species were highest in *West Bank* site, probably as a result of higher protection. Among these three recorded hornbill species; the encounter rate of Great Pied Hornbill was high in *West Bank* which was more undisturbed habitat than that of other two sites. Encounter rates of Oriental Pied Hornbill were higher in *West bank* and *Khari*, having the riverine habitat with secondary forest patches. Encounter rate of Wreathed Hornbill was almost similar in all three sites. This is probably because of their high selection range of habitat for food and roosting tree. Although the survey effort is significantly less, the encounter rate calculated from the data is much higher than that depicted by Datta (1998)^34^ i.e 1.16±0.41/ km in unlogged forest and 0.17±0.12, 0.15±0.08 and 0.10±0.04 in semi disturbed, old logged and logged forests of Pakke Tiger reserve for Great Pied Hornbill, and zero encounter of Oriental Pied Hornbill in Unlogged forest, old logged and logged forest. Encounter rate of Wreathed Hornbill not varies between the study except in the West bank region^34^. Previous studies show density of Great Pied Hornbill in Pakke varies from 3-12 individuals per sq. km ^36^ comparable with 3-10 individuals per km^2^ in western ghats^55^. Wreathed Hornbill density was estimated to be 5-15 per sq. km in 2008 in Pakke Tiger reserve^36^ was comparable with studies from Sabah 7.5 individuals per square km^56^ and 10-46 individuals per square km in East Kalimantan by Leighton 1982^57^. Study by Naniwadekar et al 2015^6^ in Namdhapa National Park also reveals that hornbill encounter rate varies between less disturbed and heavily disturbed sites.

Seed dispersers like hornbills are mutualistic to tree species assemblage in a larger landscape as they can travel a long distance tracking the food sources ^6^. But the large-scale degradation of the connectivity of protected forest area will directly affect the hornbill population, as they need undisturbed and diverse forest patches for communal roosting and nesting success ^58, 6^. The dispersal limitation of large seeded tree species also affects the diversity of tree community and will disturb the regeneration and dynamics of forest ecosystem. Comparative low abundance of hornbill dispersed tree species in Lanka was well evidenced from the present study and low abundance of hornbill dispersed species and fruiting and nesting tree denotes the change in vegetation composition in Khari and Lanka from the West bank site. Long term monitoring of the vegetation dynamics is also important for generating information on the change in vegetation dynamics in absence or limited ornithochory. The general awareness of the importance of hornbill was quite good and supportive in and around the Pakke Tiger Reserve (Personal observation) and as a result the hunting pressure is not so severe in *Lanka*. Hornbill used to move long distances in search of fruits and frequently move from undisturbed and dense forest to disturbed and logged forest ^42, 59^. Protection of vast forest patches to keep the diversity and density of the tree species intact are crucial for the survival and distribution of the hornbills and vice-versa in the landscape.

## Acknowledgements

Authors like to acknowledge Mr. Tana Tapi DFO, Pakke Tiger reserve and Mr. P. B. Rana, Range officer, Seijosa Range for their logistic support during the study period. Mr. Arun Saikia, Mr. Deba Mushahari for their help during the field work. A sincere thanks to Mr. Vivek Menon for the EDDG grant from Wildlife Trust of India to support the study. Authors also like to thank the members of Ghora Abe Society and the field staffs of Pakke Tiger reserve for their help and support during the survey period. Authors thank two anonymous reviewers for their comment that helped to improve the manuscript.

